# Benchmarking feature quality assurance strategies for non-targeted metabolomics

**DOI:** 10.1101/2021.09.09.459600

**Authors:** Yasin El Abiead, Maximilian Milford, Harald Schoeny, Mate Rusz, Reza M Salek, Gunda Koellensperger

## Abstract

Automated data pre-processing (DPP) forms the basis of any liquid chromatography-high resolution mass spectrometry-driven non-targeted metabolomics experiment. However, current strategies for quality control of this important step have rarely been investigated or even discussed. We exemplified how reliable benchmark peak lists could be generated for eleven publicly available datasets acquired across different instrumental platforms. Moreover, we demonstrated how these benchmarks can be utilized to derive performance metrics for DPP and tested whether these metrics can be generalized for entire datasets. Relying on this principle, we cross-validated different strategies for quality assurance of DPP, including manual parameter adjustment, variance of replicate injection-based metrics, unsupervised clustering performance, automated parameter optimization, and deep learning-based classification of chromatographic peaks. Overall, we want to highlight the importance of assessing DPP performance on a regular basis.

---

Liquid chromatography – high-resolution mass spectrometry (LC-HRMS) is the most widely used tool for advancing research in non-targeted metabolomics, lipidomics, and exposomics. A crucial step in hypothesis-free extraction of information from complex LC-HRMS data is automated non-targeted data pre-processing (NPP). NPP typically consists of a cascade of steps, including chromatogram extraction, chromatographic peak picking, peak alignment, and aligned feature processing (e.g., gap-filling) leading to so-called feature-tables (FT; Figure S1), which typically form the basis of any subsequent interpretation. Over the past years, numerous algorithms and software packages enabling NPP have been published (e.g., XCMS^1^, XCMS-online^2^, MZmine 2^3^, MS-DIAL^4^, El-MAVEN^5^, OpenMS^6^, and others^7,8^). Reproducing results on different tools and optimizing parameters has been recognized as one of the main challenges in non-targeted data processing, impacting data interpretation and ultimately study outcome.^9–15^ The emerging unease regarding the under-utilization of data^12,13^, calls for evaluation strategies that enable benchmarking of different tools, algorithms, and parameter choices on a regular basis. Undeniably, validation of LC-HRMS NPP poses several challenges. First, the true number of peaks (and their properties such as area, height, etc.) in a raw dataset is typically unknown. This is even true when measuring a defined set of LC-HRMS grade analytical standards due to unavoidable chemical impurities, contaminations, unexpected adducts, and gradient artifacts. Second, no accepted gold-standard NPP method which would allow the generation of reliable reference FT as benchmarks exists. Finally, datasets differ in their characteristics (peak widths, mass precision, etc.), which implies that conclusions on NPP performance drawn from a benchmark might only allow conclusions for other datasets with similar characteristics. State-of-the-art NPP reliability assessment strategies include peak/feature classification^16–18^, unsupervised clustering^18–21^, and benchmark recovery-based^9,22,23^ techniques, with the first two being applied almost routinely, while recovery-based approaches were mainly limited to dedicated NPP assessment studies. Up to date, there is no standardized catalog of metrics enabling the assessment and reporting of general figures of merit. In fact, the opposite is the case: there is vast heterogeneity in used metrics and their ability to describe NPP performance.

Figure 1 provides an overview of current classification-based NPP assessment strategies, their metrics, and general capabilities. Generally, classification discerns reliable from unreliable peaks/features (e.g., reliable LC-HRMS peaks should co-elute with isotopologues identifiable via specific mz distances), and many principles can be easily implemented as NPP performance tests. However, key aspects of NPP quality remain blind spots. First, for peaks/features classified as unreliable, it is generally not possible to differentiate between cases where chromatographic noise has been erroneously picked as peak and cases where chromatographic peaks have been picked poorly. Moreover, the majority of principles allowing to assess the quality of reported abundances yields a checkpoint only after alignment. Therefore, it is difficult to trace the problems back to the peak picking or alignment step. Finally, classification-based methods do not assess the proportion of undetected peaks (false negatives).

**Figure 1.**
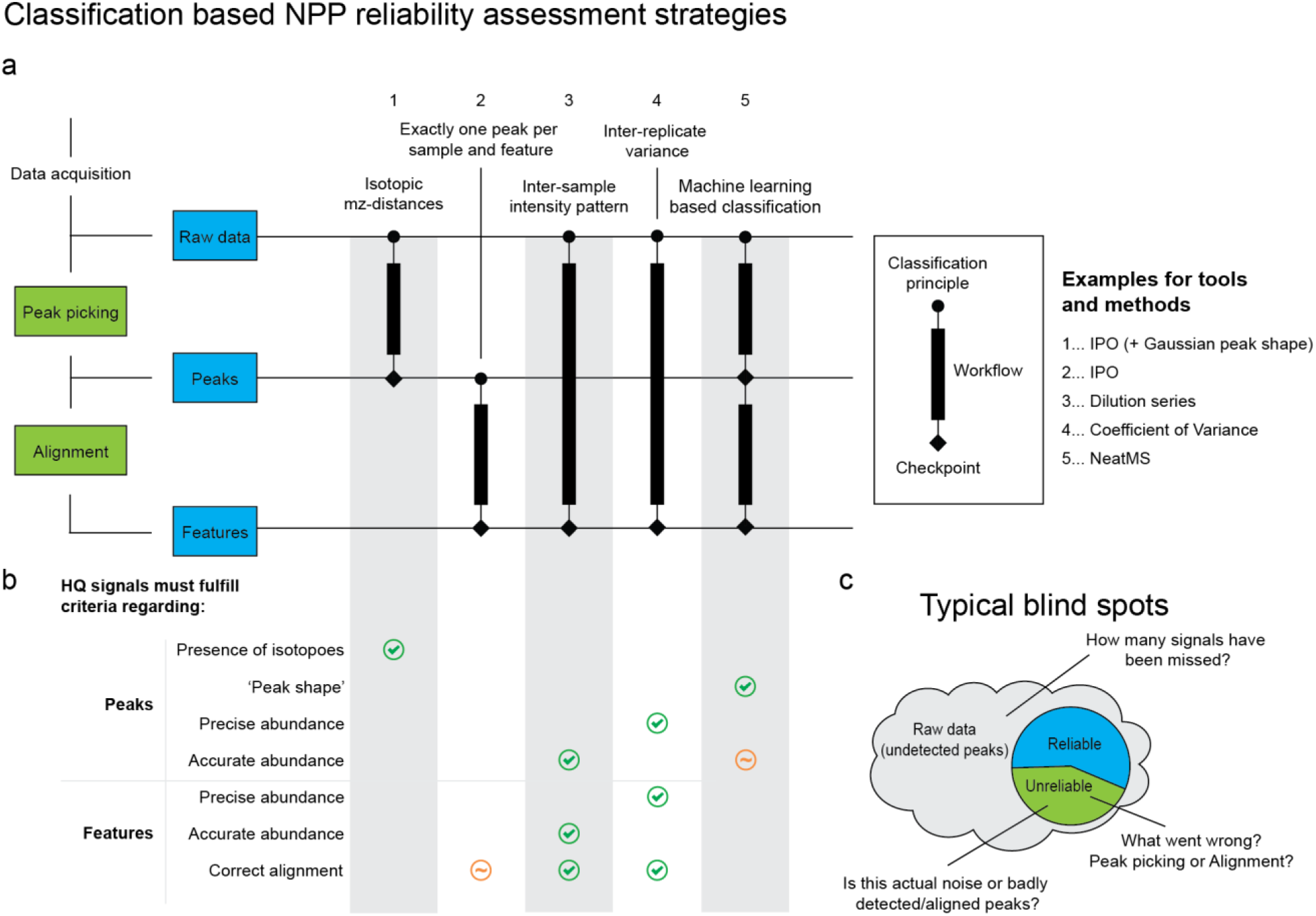
Selected strategies for classification-based performance assessment for non-targeted data pre-processing (NPP) of LC-HRMS data. (a) Five principles distinguishing reliable high quality (HQ) from unreliable peaks/features have been summarized. While some approaches require only peak picking for their application, others can only be utilized after alignment. (b) Different principles allow for the assessment of data properties. Notably, in cases where multiple criteria have to be fulfilled, all of them have to pass for a peak/feature to be considered HQ. (c) Typically, classification-based approaches suffer from a number of blind spots.

NPP assessment through comparison with targeted data evaluation, denoted as benchmark recovery-based approaches, has the potential to investigate these blind spots. While the concept is straightforward, it is rarely applied in routine non-targeted experiments, as its implementation can be tedious. This is because benchmark generation requires meticulous manual curation of peaks, which can be very time-consuming and is often considered to be too subjective for ground-truth generation. In fact, a recent study showed that three experts in mass spectrometry strongly disagreed on what constitutes an actual chromatographic peak in ∼20% of cases (n = 1071)^24^, demonstrating how vague boundaries between differences in opinion carry the risk of overinterpreting differences between benchmarks and NPP results. Moreover, the manual work of benchmark curation leads to rather small sets of benchmark peak lists, potentially hampering the representativeness of the benchmarks for the whole dataset.^14,21^

We recently introduced mzRAPP, a tool that allows simplifying benchmark recovery-based NPP assessment while also increasing its reliability and providing functionality for their utilization.^25^ Briefly, it takes the outputs of traditional targeted metabolomics data evaluation (molecular formulas with retention time boundaries per sample) as an input and automatically increases the number and reliability of benchmark peaks via consideration of isotopologues. In this way, benchmark peak abundances can be confirmed by predicted isotopologue ratios (IRs), rather than solely by human judgment. Finally, the consideration of isotopic relations allows the introduction of performance metrics dissecting the individual steps of NPP (e.g., see Figure S2).

NPP results obtained following the steps of peak-picking, alignment, and gap-filling, respectively are benchmarked using metrics such as the recovery of benchmark peaks and isotope ratio biases (IRbs). The latter reflects the accuracy of the peak abundances obtained as visualized in Figure Figure S2a. As another key advantage, metrics on the NPP alignment process can be retrieved without requiring tedious curation of the correct alignment of all benchmark peaks. By simply counting cases where the alignment performed on a given isotopologue is in conflict with the alignment of another isotopologue of the same molecular species (see Figure S2b). Building upon these principles, mzRAPP generates benchmarks and performs reliability NPP assessment at different stages of the process (picked peaks, peak alignment, and processed features). Currently, mzRAPP supports outputs from more than five of the most commonly used NPP tools (XCMS, XCMS-online, MZmine 2, MS-DIAL, OpenMS, El-MAVEN, MetaboanalystR 3.0^26^, etc.).

In this work, we show the power and necessity of this novel validation scheme. We scrutinize current NPP execution and assessment strategies by adding a new validation layer as enabled by representative high-quality benchmarks created case by case. Moreover, we explore the most common NPP optimization/assessment strategies, including the most recent NPP classification approaches based on artificial intelligence, by showing their capability and limitations in an unprecedented manner. For the first time, cross-validation, including the quantitative dimension of features/peaks, is within reach in the otherwise black-box-like environment of NPP.

## Methods

### Datasets

Datasets used for benchmark generation were downloaded from Metabolights^27^ or thankfully provided by the authors of the respective studies.^9,22,24,28–31^ References and/or repository IDs are provided in column ‘Reference’ of Table S1. Where not already provided as centroided mzML files, raw files were centroided and converted to mzML format via ProteoWizards msConvert^32^ (version 3.0.21045-7732b6429). Names of raw files used are provided in Table S2. Files acquired via fast polarity switching were filtered to contain only positive scans.

### Benchmark generation

Targeted data evaluation was performed via Skyline^33^ (version 21.1.0.146) and for the most abundant isotopologue of each molecule, for which retention time and molecular formula were known, in all datasets. Then, manual set retention time boundaries were exported for each molecule and mzML file. These and the mzML files formed the input for mzRAPP, which also extracted other predictable isotopologues for each molecular formula. Only isotopologues with a Pearson Correlation Coefficient (PCC) > 0.85, below an isotopologue ratio bias (as calculated by peak areas) of 35%, an isotopologue ratio bias (as calculated by peak heights) of 30%, and a difference between ratio bias (height) and ratio bias (area) below 30% points were accepted.

### Extraction of non-targeted data pre-processing performance metrics

Extraction of NPP performance metrics was conducted via mzRAPP (version 1.1.6). Exact criteria and rules for matching between signals of the benchmark and those of the unaligned and aligned NPP outputs can be found in the original mzRAPP publication^25^ and on Github (https://github.com/YasinEl/mzRAPP). Isotopologue ratio biases (IRbs), as calculated from NPP outputs, were considered to be recovered if they were less than 20 %points higher than respective BM IRbs. Confidence intervals (confidence level = 0.99), for all NPP metrics were derived via bootstrapping of benchmark molecules (R = 1000) using the boot package (version 1.3-28).

### Application of non-targeted data pre-processing

All data pre-processing experiments were performed via XCMS3 (version 3.14.1) using R 4.1.0 and MZmine 2 (version 2.53). Parameter optimizations were performed manually or via automated optimization tools. The automated optimization tools were IPO^17^ (version 1.18.0), AutoTuner^34^ (version 1.6.0), MetaboanalystR 3.0^26^, and SLAW^35^ (version 1.0.0). Classification of peaks of peaks by quality was performed via NeatMS version (0.9), which was run via Python 3.7. For the parameter sensitivity study, the coefficient of variance investigation, and the parameter optimization dataset (DS) 5 was processed. For the unsupervised clustering investigation, and the assessment of NeatMS DS 1 was processed. Additional details on set parameters for all performed studies are given in the supplementary material.

### Data analysis and figures

All further data analysis was performed using R (version 4.1.0) and R studio (version 1.4.1717) using the data.table package. Plots were generated using ggplot2, patchwork, and ggradar. Figures and diagrams were further processed using Adobe Illustrator (25.3.1).

## Results and Discussion

### The quality of automated benchmark curation and extension

Ideally, a BM used for NPP assessment should be produced from the same dataset or a dataset generated via the same instrumental platform as the dataset of interest. In this study, BMs from 11 different public and in-house raw datasets (listed in Table S1) were generated via mzRAPP. The case by case generated BMs covered five different MS-systems, coupled to hydrophilic interaction chromatography (HILIC) or reversed-phase chromatography (RPC), as well as different sample types (including analytical standard mixtures, blood serum, red blood cell extracts, and cell culture extracts) and compound classes (polar metabolites, lipids, and exogenous small molecules). Targeted extraction of the most abundant isotopologue of each known molecule was done manually, but was automatically extended to all lower-abundance isotopologues. Quantitative properties of the thereby generated benchmarks are visualized in Figure S3. In Figure S3a all 46742 BM peak areas of low abundant isotopologues (LAITs) were plotted against the area predicted from the respective most abundant isotopologue (MAIT), showing excellent linearity. It should be noted that only peaks within the linear dynamic range of the instrumental platform were added to the BMs. Figure S3b shows the absolute peak area bias of all LAITs, with 94% of all calculated isotopologue ratio biases < 25%. Comparison of biases (Figure S3c) as calculated via peak areas versus peak heights (which are generally more robust as they are not affected by poor setting of RT boundaries) revealed good agreement, further strengthening the evidence of accurate “ground-truth” for an extensive number of peaks. Finally, for DS1, the benchmark reliability was evaluated upon comparison with reported fold changes assessed in an independent laboratory. Figure S3d shows the excellent reproducibility for the mzRAPP approach applied here. For this specific dataset, the number of peaks with reliable quantitative properties increased by > 200% by integrating LAITs. The addition of LAITs increased not only the BM size but also the covered dynamic range. Figure S4a and b quantifies this significant extension for all eleven investigated datasets.

Peak metrics such as the full width half maximum (FWHM) of chromatographic peaks and the mass precision given by mz ranges of individual peaks are important for any dataset to be analyzed via NPP. In fact, most NPP tools require parameters corresponding to these variables to be set for any dataset to be processed. Therefore, it is worth noting that generated BMs showed large differences in all these metrics as a result of different measurement methods (see Figure S4c and d). Next to these characteristics, the peak shape as reflected by the zigzag index^39^, sharpness^40^, and other metrics (Figure S5) show significant differences across investigated datasets. This highlights the importance of using benchmarks very similar to the dataset of interest (and at best generated from the very same dataset) for potential NPP assessment studies.

Ultimately, benchmarks can be utilized to derive performance metrics for NPP experiments conducted on the same raw datasets. As outlined above, mzRAPP enables automated assessment of the proportion of recovered benchmark peaks and the proportion of recovered isotopologue ratio biases (IRbs) before and after alignment as key NPP performance metrics. Moreover, other metrics such as the proportion of NPP peaks with reported integration boundaries close to the intensity maximum of a benchmark peak are reported.

Finally, the metrics derived from the benchmark should be translatable into an actual proportion for the underlying dataset. Representative sampling of the benchmark peaks/features is a prerequisite. To show the validity of our approach, we estimated confidence intervals (CI) for all metrics by bootstrapping benchmark molecules. In Figure 2 and Figure S6 we show how the number of molecules used for BM generation affects metrics derived from NPP reliability assessment. This was done by bootstrapping different numbers of molecules from BM 1 (containing 712 molecules and > 30000 peaks). It can be observed how a reduction of the number of BM molecules increases the CI while the assessed metric was in agreement in almost all cases. Therefore, even < 100 BM molecules can lead to reasonable estimates of the performance of NPP, as long as the increase in CI can be accepted.

**Figure 2.**
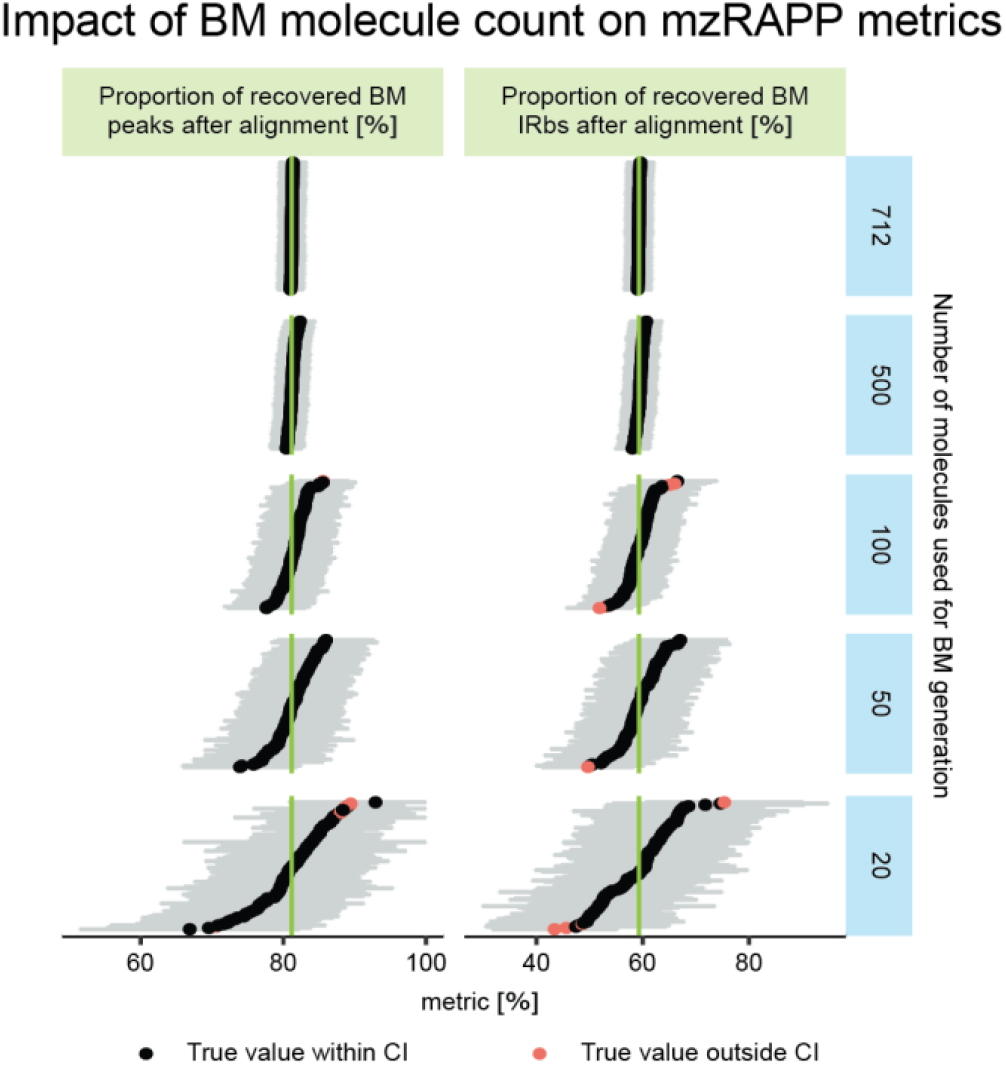
A benchmark was generated for dataset 1 considering 712 molecules and utilized for NPP assessment. Stepwise reduction of molecules used for BM generation (molecules were sampled at random from all 712 molecules (n = 100) for molecule numbers ranging from 20 to 712) largely affected the size of confidence intervals (CI; as estimated by mzRAPP via bootstrapping with R = 1000 and confidence level = 0.99), while true metrics, as estimated from the original BM (green lines) stayed largely within within the reported CI of respective NPP assessments. (CI values above 100 were reduced to 100)

### Sensitivity of NPP parameters

NPP requires the adaptation of different parameters to the analyzed dataset. These parameters can appear more or less intuitive to users with different scientific backgrounds and experiences. Generally, parameters involving expected chromatographic peak widths and retention time shifts are often considered to be among the more intuitive parameters. In the following, we showcase examples that emphasize the need for case-by-case benchmarking strategies, as even intuitive parameter settings in the NPP could have an adverse impact on the NPP output.

Figure 3a shows how a stepwise increase of XCMS’s cent-wave’s maximum peak width parameter (MPW) using 2 s increments heavily affected the proportion of missed BM peaks and increased BM isotopologue ratio biases (IRbs). In the most extreme case, an increase in MPW from 26 to 28 s led to an increase in the proportion of missed BM peaks (before alignment) from 6 to 93%. Considering that the median of BM peaks full-width half maxima (FWHM) ranged from 4 to 18 s with a median of ∼7 s, there was no trivial dependence of the optimal MPW on the FWHM of peaks to be detected. While fewer peaks were missed after alignment and gap-filling, this improvement was insufficient to make up for errors introduced during peak detection. It is worth noting that even in cases where gap filling recovered most peaks, such as with an MPW of 14 s, the resulting peak areas led to worse IRbs than when peaks were already detected in the peak detection step (e.g., with MPW set to 12 s). While the highest observed retention time shift in the BM peaks was below 10 s, the maximum allowed shift, as set via the band-width (bw) parameter in the group density algorithm, did not affect NPP results to the same extent as MPW. Interestingly, there was almost no effect of the set MPW on IRbs after peak picking. However, there was a significant impact on IRbs after peak alignment and gap-filling, which depended on the MPW set during the peak picking step rather than set alignment parameters.

**Figure 3.**
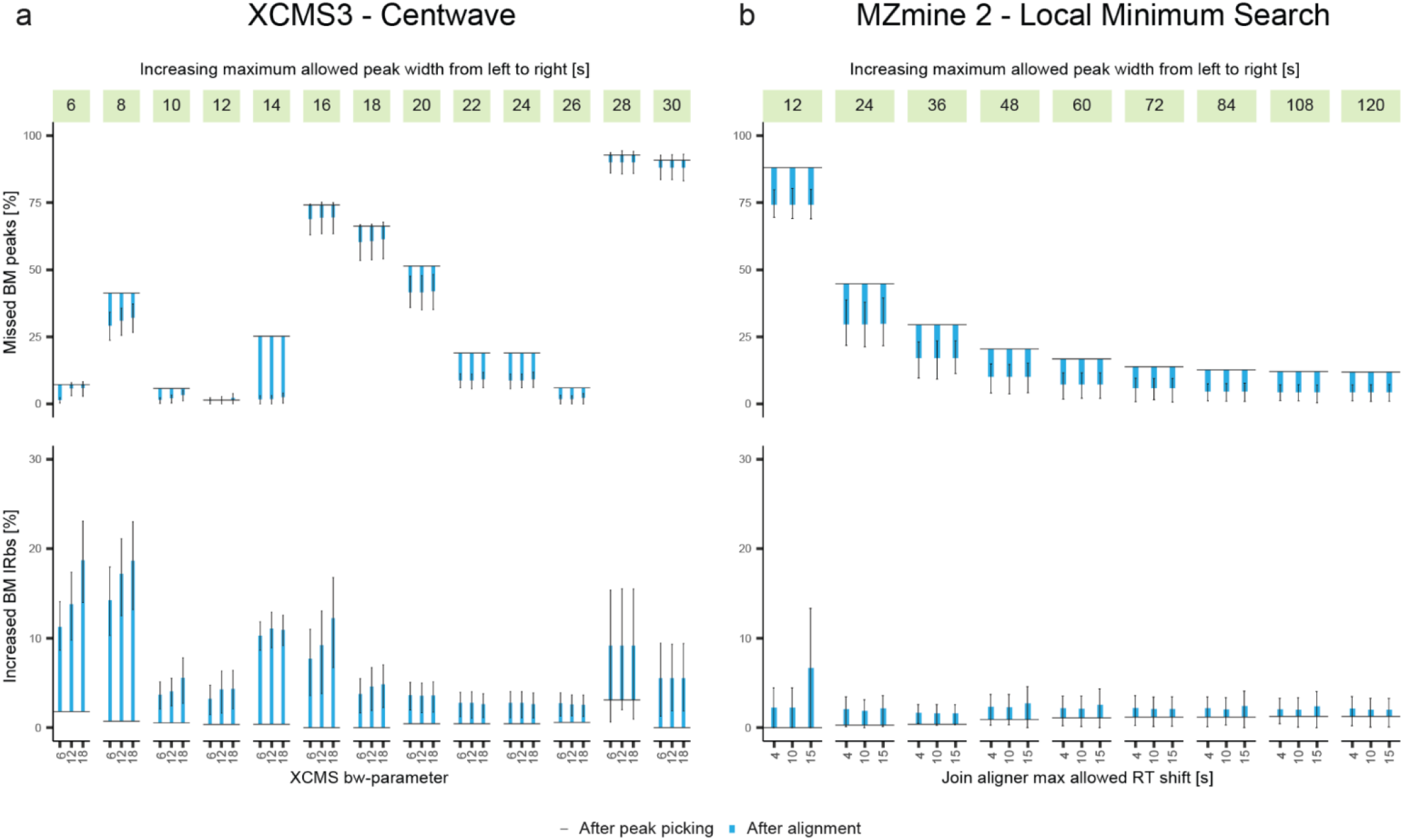
Multiple non-targeted data pre-processing experiments with parameter sweeps for the peak-picking and the alignment step were performed on the same dataset via XCMS3 and MZmine 2. The peak picking parameter referring to the maximum peak width allowed (MPW) was incrementally increased for both tools, with three retention time tolerances tested for each peak picking step. (a) The proportion of recovered BM peaks was strongly and often abruptly affected by the MPW parameter. While the proportion of increased isotopologue ratio biases (IRbs) was more strongly influenced after alignment, this effect was primarily dependent on the MPW parameter during peak picking. (b) In MZmine 2, an increase of the MPW never led to a reduction in the proportion of recovered BM peaks.

The same dataset was processed via MZmine 2 and incremental increase of the MPW using its Local Minimum Search peak detection algorithm as visualized in Figure 3b. Here, the relationship between the set MPW and the proportion of recovered peaks was found to be as expected, with more BM peaks being recovered with increasing MPW. However, the absolute MPW necessary to reach the optimum of ∼20% missing BM peaks after peak detection was > 72 s, a factor of ∼10 higher than the median BM FWHM of ∼7 s. While this algorithm assessed the MPW at the base of peaks rather than at half maximum, this might still lead to problems since the base width of a chromatographic peak can vary widely for real-world applications. The IRbs were almost unaffected by the MPW, with only one outlier for an MPW of 12 s and a retention time tolerance of 15 s.

For this specific data set, the optimum of all parameter sets tested was found by XCMS (MPW of 12 s and retention time tolerance of 6 s, leading to a proportion of missed BM peaks < 1% and IRbs < 5%). It should be noted that this finding cannot be generalized, but it holds true for the processed data set, representing a use case of parameter adjustment. In fact, the conclusion that MZmine 2 is generally underperforming as compared to XCMS would be wrong since testing the entire parameter space was beyond the scope of this study. The example clarifies that the common practice of manual parameter adjustment to metrics derived from the analytical performance of the instrumental setup can lead to suboptimal NPP results.

### Application of NPP parameter optimization tools

Current NPP parameter optimization tools (as implemented in IPO^17^, AutoTuner^34^, MetaboanalystR 3.0^26^, and SLAW^35^) undoubtedly facilitate the NPP step in non-targeted analysis and improve quality. Here we test the classification-based tools applying our benchmark-recovery approach. This way, the otherwise missing metrics of missing peaks and accuracy of peak abundances are validated. Figure 4 compares the quality of NPP-parameter optimization performed for the NPP of DS 5 via different metrics as exported by mzRAPP. As can be seen, the differences between the optimization attempts were observable for the proportion of missed benchmark peaks and increased IRbs before and after alignment, as well as the proportion of BM peaks leading to split peaks. It turned out that the initially defined values for the parameter optimization process is crucial and was unique for each tool. For example, in the case of IPO, manual adjustment of starting parameters led to a decrease in the proportion of BM peaks missing after alignment from ∼25% to ∼1%. Again, this test emphasizes that NPP evaluation is not redundant when using automated parameter optimization algorithms.

**Figure 4.**
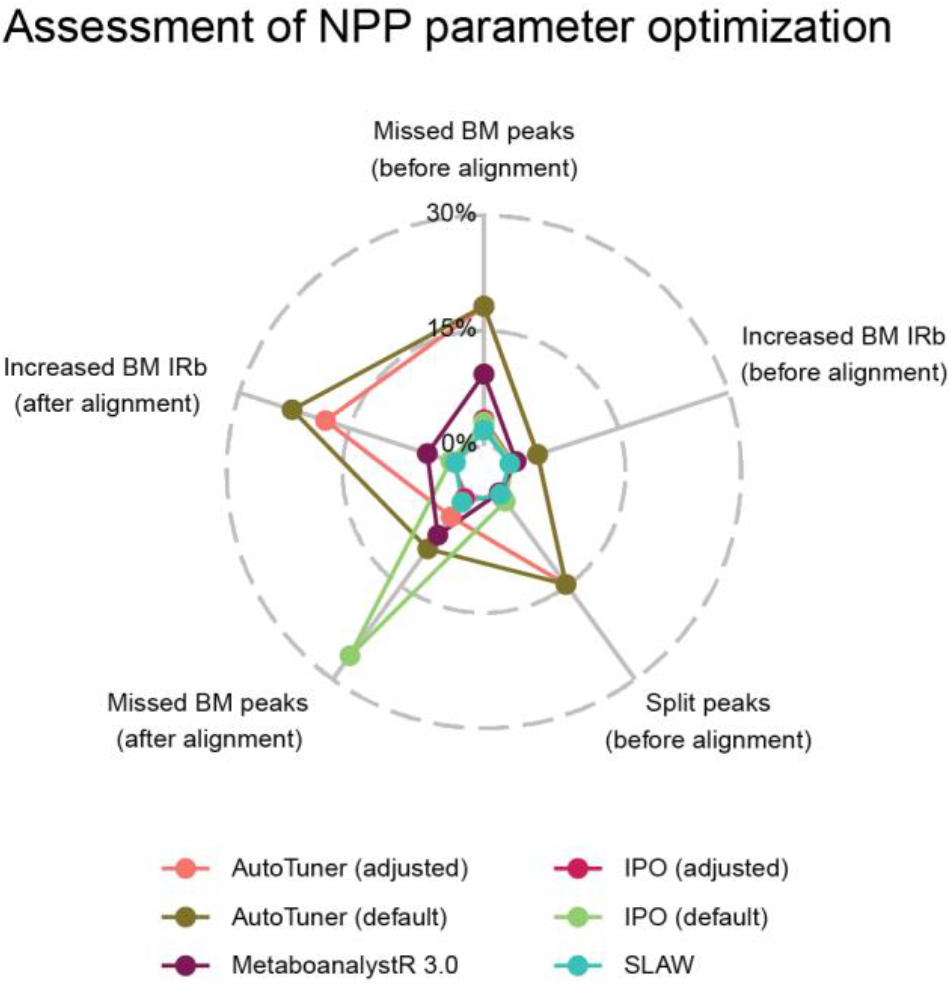
Non-targeted data pre-processing (NPP) parameters for the processing of dataset 5 have been optimized using different optimization tools (IPO and AutoTuner, both adjusted and default, as well as MetaboanalystR 3.0 and SLAW). Outcomes were assessed via five benchmark recovery-based metrics, namely the proportion of missed benchmark (BM) peaks (before and after alignment), the proportion of increased isotopologue ratio biases (before and after alignment), and the proportion of split peaks (before alignment). Confidence intervals of all metrics for the underlying dataset are given in Table S3.

### NPP assessment or filtering via Coefficient of Variance

A widely accepted metric in NPP assessment is the Coefficient of Variance (CV) obtained from replicate measurements. It is common practice to report the number or proportion of NPP features with a Coefficient of Variance (CV) below a certain threshold (commonly 30%). In Figure 5, this otherwise straight-forward approach is investigated. CV-based quality metrics are plotted versus proportions of recovered BM peaks and IRbs for a total of 78 NPP experiments (using different values for the maximum allowed peak width in the peak-picking step and different values for bw in the peak alignment step) performed on DS 5. The number and proportion of features with a CV < 30% (nCV30 and pCV30, respectively) were assessed. For a given bw setting, the metric nCV30 was well correlated with the proportion of BM peaks recovered post-alignment (e.g., PCC = 0.9 for bw = 0.5), while the proportion of features with CV < 30% reflected the proportion of IRb recovered post-alignment (e.g., PCC = 0.97 for bw = 0.5). Filtering NPP features to remove features with missing peaks and setting a threshold of CV > 30% led to an increase in the proportion of recovered IRbs (post-alignment) in all cases. In contrast, the proportion of overall recovered peaks decreased. Thus, the trends in nCV30 were indicative of the proportion of found peaks, while the trends in pCV30 corresponded to the proportion of high-quality peaks. The application of this filter successfully removed low-quality features contributing to increased IRbs. This highlights how classification-based NPP assessment approaches do not distinguish between noise and badly detected peaks, as discussed above (as well as in Figure 1). It should be noted that the high correlation of pCV30 with the proportion of recovered IRb (post-alignment) persisted only as long as the bw parameter was kept constant. For example, while the trends of pCV30 were very similar in all settings of bw, the proportion of recovered IRb (post-alignment) for bw = 0.5 decreased for all MPW values. As a consequence, pCV30 is not a suitable metric to evaluate whether one given NPP result has higher quality than another. The same is true for nCV30. In fact, the highest nCV30 (for bw = 0.5) would have led to ∼75% recovered IRbs (after alignment), which is far from the optimum of approximately 100% (with values > 2 or 18). Thus, both CV metrics have their valid role in quality assessment. However, simple NPP optimization based on CV metrics is precluded. Neither optimizing nCV30 nor the pCV30 led to a complete recovery of BM peaks and BM IRb and thus to an optimum result.

**Figure 5.**
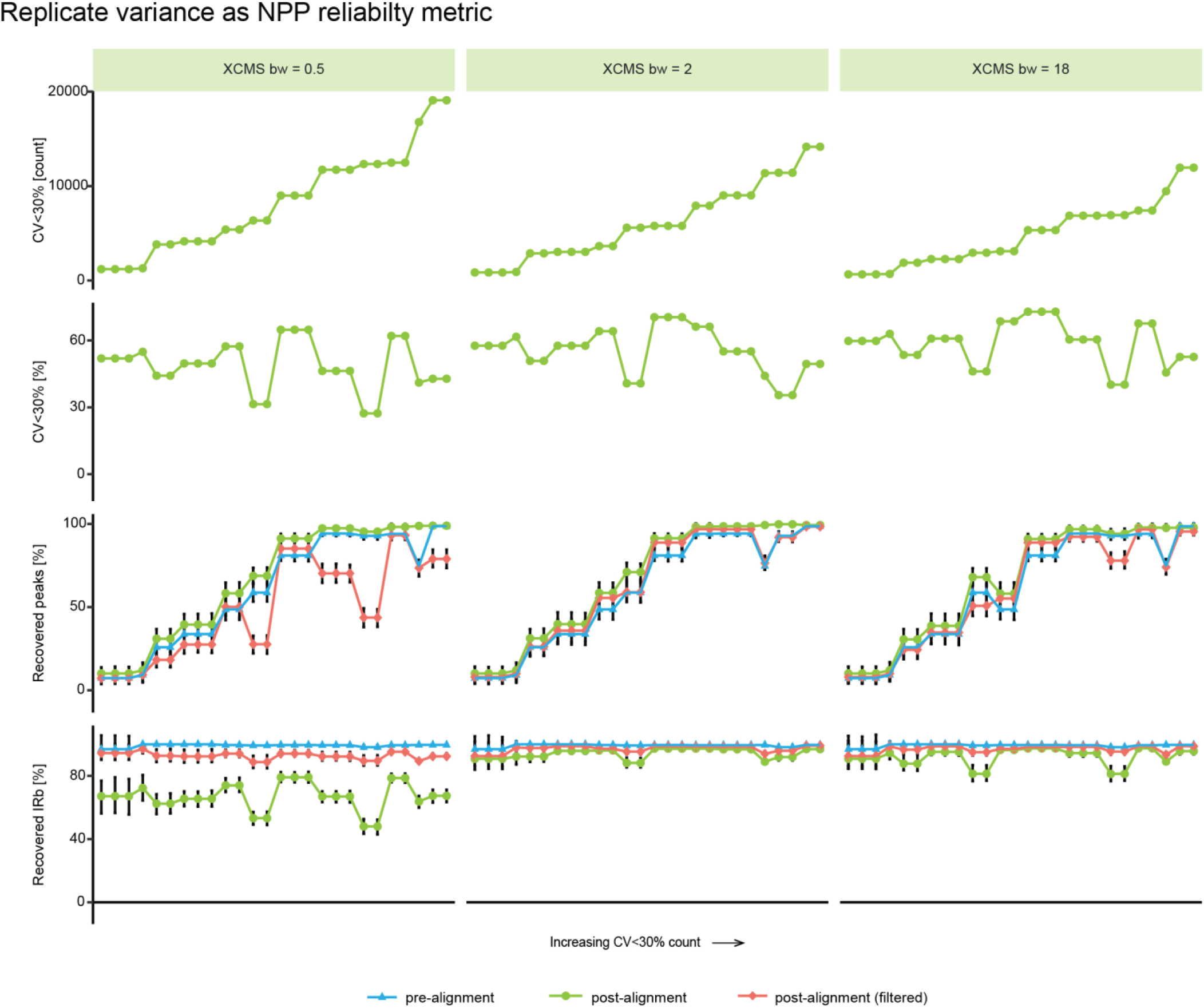
A dataset consisting of 9 replicate injections was processed via XCMS using different values for the maximum allowed peak width (MPW) parameter and the bandwidth (bw) parameter, leading to a total of 78 non-targeted data pre-processing experiments. Four different quality metrics including the number of features with CV < 30%, the proportion of features with CV < 30%, and the proportion of recovered benchmark peaks and isotopologue ratio biases (IRb) (pre-alignment, post-alignment, and post-filtering (only features without missing values and with CV < 30%)) were then plotted (sorted by increasing number of features with CV < 30%).

### NPP assessment via unsupervised clustering

Unsupervised clustering approaches such as Principal Component Analysis (PCA) are an integral part of non-targeted workflows and their role in discovery is undisputed. However, they are not suitable for NPP-quality assessment, as emphasized in Figure S7 and Figure S8. PCAs performed on the NPP outputs showed good separation of sample groups, not reflecting the benchmark recovery metrics, and thus not validating the reliability of the NPP output.

### Application of peak classification via artificial intelligence

Novel tools such as NeatMS use artificial intelligence for the classification of NPP peaks by their quality. As a major breakthrough, noise removal is accomplished without relying on replicate injections. Successful application of machine learning algorithms requires good training data, which (in the case of NeatMS) have to be labeled by users with different skill sets. In this work, we scrutinized NeatMS. NPP peaks generated from nine NPP experiments performed on the same dataset (DS 1; containing 10 samples), were classified accordingly, into three categories ‘High quality’, ‘Low quality’ and ‘Noise’. We then applied different filters to the aligned NPP features and required them to contain 0, 1, 3, 5, 8, or 10 ‘High quality’ peaks. Subsequently, the proportion of recovered peaks and IRbs after alignment was assessed. For this purpose, we filtered our BM to contain only features with peaks in all 10 samples. As can be seen in Figure 6, removing all NPP features which did not include at least 1 ‘High quality’ peak reduced the number of NPP features by ∼40 to ∼60%, while having almost no effect on the proportion of recovered BM peaks or IRbs, demonstrating how NeatMS can be applied successfully for removing false positives from NPP results. However, requiring more ‘High quality’ peaks reduced the proportion of recovered BM peaks by multiple %points in many cases. When all 10 samples were required to contain only ‘High quality’ peaks for a feature to be retained, the proportion of recovered BM peaks dropped <10% in all cases. Our validation confirms that tools such as NeatMS for efficiently removing false-positive signals from NPP results significantly advance NPP. Despite this undisputed role, the quality and size of training data strongly affects the procedure and is defined by a user, case by case. Thus, independent validation continues to be of great value.

**Figure 6.**
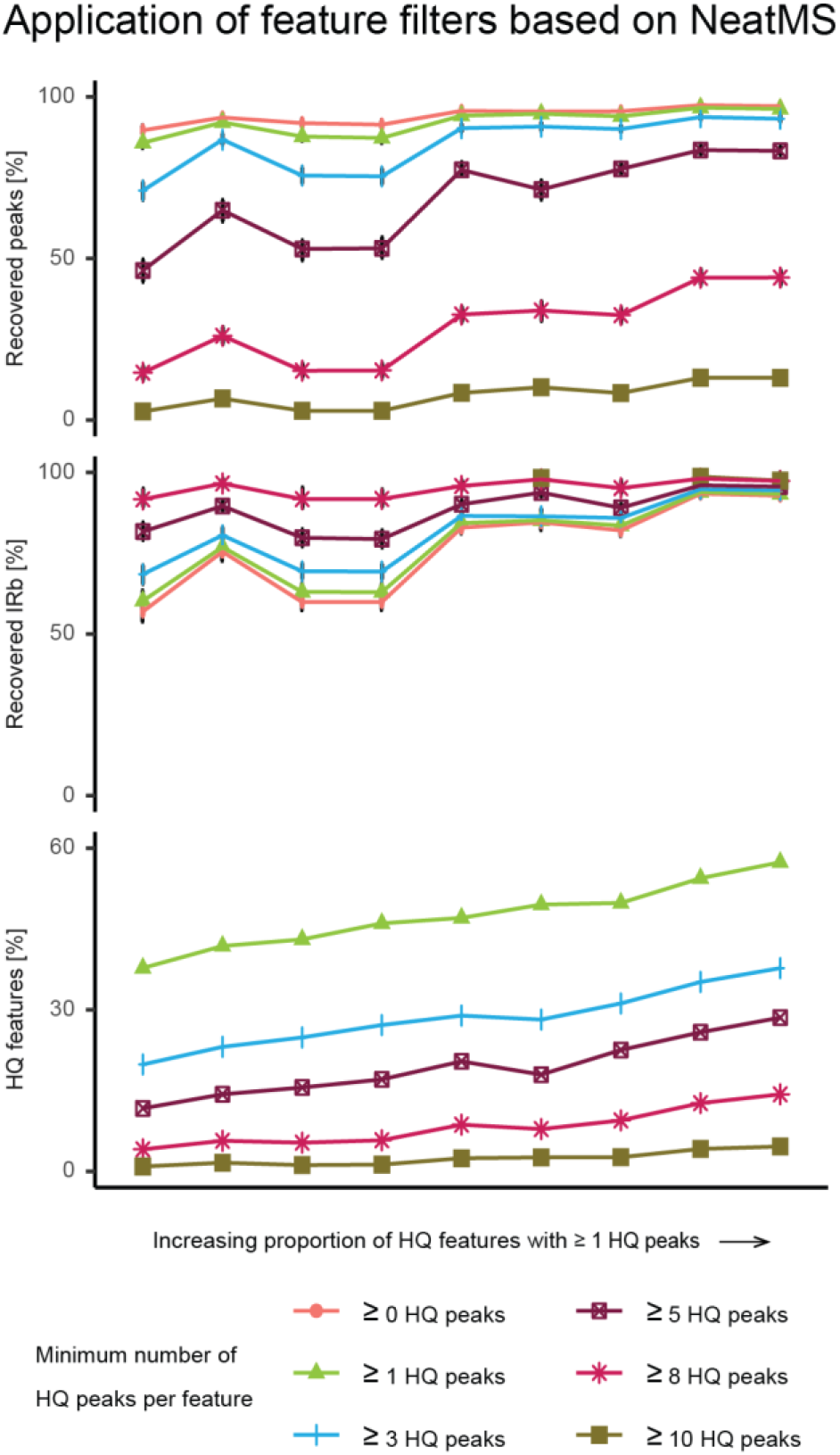
Dataset 1, consistent of 10 samples, was processed with 9 different sets of XCMS-parameters. All peaks produced via XCMS were classified by NeatMS into different categories, including high quality (HQ), or noise. Different numbers of peaks within an aligned feature were required to be of HQ for a feature to be retained. The plot on the bottom shows the proportion of all XCMS features satisfying these criteria for each parameter set. The plots above show metrics on the proportions of recovered benchmark peaks and recovered isotopologue ratio biases (IRbs). The x axis was sorted by increasing values of HQ features [%] for more than or equal to 1 HQ peaks.

## Conclusion

The study emphasized the need of case-by-case optimization and validation of NPP. Due to the described obstacles in manually adjusting parameters, even when applying cutting-edge optimization tools, we reason that NPP performance should be evaluated on a regular basis. Moreover, discussed problems in finding NPP optima via strategies such as unsupervised clustering or Coefficient of Variance-based metrics entail the necessity for alternative assessment methods. NPP can perform in unpredictable ways, and outputs should be assessed against a solid ground-truth which cannot be generalized in the form of a golden dataset, against which all NPP algorithms are optimized. We demonstrated how case-by-case benchmark recovery-based approaches can satisfy this need. The fact that benchmark information can be based on a solid foundation of orthogonal information allows for increased reliability and accuracy. mzRAPP can produce representative and reliable benchmarks for each investigated dataset, which can be integrated for routine validation of non-targeted NPP. Finally, the performance ranking of different NPP tools (as reported elsewhere) is not useful.

## ASSOCIATED CONTENT

### Supporting Information

Supplemental Methods, Figures S1 – S8, and Table S1 (PDF)

Table S2 (CSV)

Table S3 (CSV)

## AUTHOR INFORMATION

### Author Contributions

The manuscript was written through contributions of all authors. / All authors have given approval to the final version of the manuscript.

### Notes

The authors declare no conflict of interest. Where authors are identified as personnel of the International Agency for Research on Cancer/World Health Organization, the authors alone are responsible for the views expressed in this article, and they do not necessarily represent the decisions, policy or views of the International Agency for Research on Cancer/World Health Organization. R.S. would like to acknowledge support from ANR-19-CE45-0021 (MetClassNet) grant.

## ACKNOWLEDGMENT

The authors would like to acknowledge Yoann Gloaguen for his advice regarding the application of NeatMS.

