## Supplemental Methods, Figures S1 - S8, and Table S1 for "Benchmarking feature quality assurance strategies for non-targeted metabolomics"

### Table of Contents:

**Supplemental Methods.**

**Figure S1.** Major steps of non-targeted pre-processing and glossary

**Figure S2.** Benefits of isotopologues in benchmarking non-targeted pre-processing

**Figure S3.** Quality metrics for mzRAPP benchmarks

**Figure S4.** Impact of isotopologues on benchmark sizes and distributions of chromatographic FWHM and mz ranges

**Figure S5.** Different variables extracted for all benchmark peaks

**Figure S6.** Impact of number of molecules used for benchmark generation

**Figure S7.** mzRAPP metrics for nine non-targeted experiments for dataset 1 & two principal component analysis performed on the same outputs.

**Figure S8.** Principal Component Analysis performed on non-targeted outputs shown in Figure S7

**Table S1.** Overview of used datasets and benchmarks

### Supplemental Methods

#### Non-targeted data pre-processing parameter sensitivity study

DS 5, which consists of 9 replicate injections of the same sample, was processed via XCMS3 (version 3.14.1) using R 4.1.0, and MZmine 2 (version 2.53). For XCMS3 we applied the Centwave[29] peak detection algorithm with prefilter = c(3, 100), ppm = 5, noise = 5, snthresh = 3, fitgauss = TRUE, integrate = 2 and mzdiff = -0.002. In the peakwidth parameter we set the minimum allowed peak width to 1. The maximum allowed peak width was increased from 6 to 30 s with steps of 2 s. All other parameters were at their default values. Alignment of peaks was performed via the PeakDensity algorithm with minFraction = 0.1, and binSize = 0.002. For the bandwidth (bw) parameter the values 6, 12 and 18 were tested. All other parameters were left at default. Filling of gaps was done via the fillChromPeaks algorithm, with all parameters left at their default values. For MZmine 2, mass detection was performed via Mass detector = Centroid and Noise level = 0. ADAP Chromatogram builder[30] was used for chromatogram extraction with Min group size = 5, Group intensity threshold = 1000, Minimum highest intensity = 15000, and m/z tolerance set to 0.003 m/z or 5 ppm. Peak detection was performed via Local Minimum Search with a chromatographic threshold = 30%, Search minimum in RT range = 0.2, Minimum relative height = 5%, Minimum absolute height = 15000, Minimum ratio peak top/edge = 2, and Peak duration range starting from 0.05 min. The maximum allowed peak duration was screened using values from 0.2 to 2, with a step size of 0.2. Peak alignment was performed via Join aligner with m/z tolerance = 0.003 or 3 ppm, weight for m/z = 0.8, weight for RT = 0.8 and Retention time tolerance values 4, 10 and 15 s. For the extraction of performance metrics via mzRAPP, we used benchmark 5.

#### Coefficient of Variance investigation

DS 5 was processed via XCMS3 in the same manner as described for the parameter sensitivity study, but with different maximum allowed peak width and bw values. Specifically, maximum allowed peak width values were screened from 5 to 30 s with a step size of 1 s, and bw values were set to 0.5, 2, or 18. For the extraction of performance metrics via mzRAPP, we used benchmark 5, but filtered it to contain only features with a peak in each sample and a Coefficient of Variance < 30%, as calculated via chromatographic peak areas.

#### Unsupervised clustering investigation

DS 1 was processed via XCMS3 using centwave for peak detection with prefilter = c(2, 10000), ppm = 3, noise = 1000, snthresh = 20, mzdiff = 0.03, and maximum allowed peak width = 20. The minimum allowed peak width was set to values ranging from 1 to 3, with a step size of 1. Peak alignment was done via PeakDensity algorithm with minFraction = 0.1, and binSize = 0.003. For bw the values 2, 6, and 25 were tested. Extraction of performance metrics was conducted via mzRAPP, using BM 1. Principal component analysis was conducted for all aligned XCMS outputs (after scaling and centering) within R using the stats package.

**Application of parameter optimization tools**

Parameters were optimized via IPO[16] (version 1.18.0), AutoTuner[31] (version 1.6.0), MetaboanalystR 3.0[26], and SLAW (version 1.0.0) for dataset 5. For IPO parameters were optimized once from default starting values of IPO and once with slightly adapted starting values. Specifically, min_peakwidth was set to 3-8, max_peakwidth to 30-50 ppm to 1-5, noise to 0-5000, bw to 3-20, mzwid to 0.005-0.02, and minfrac to 0.1. Optimized parameters were then applied via XCMS. For AutoTuner lag was set to 22, threshold to 3.1 and influence to 0.15 for initial processing. For 'default' application of AutoTuner we only set parameters optimized by AutoTuner in XCMS3 and left the rest at the default values from XCMS3. In a second attempt we manually set the binSize of the PeakDensity algorithm in XCMS3, which is not optimized by AutoTuner, to 0.005. For optimization via MetaboanalystR 3.0 we set the starting parameters via their implemented platform selection to 'UPLC-Q/E', and set the polarity to 'positive', and the minFraction to 0.2. Pre-processing was then conducted within the MetaboanalystR 3.0 package. For SLAW we set the peak detection algorithm to 'centwave', and optimization = true. Pre-processing was then conducted within SLAW.

**Assessment of NeatMS**

NeatMS[32] was run via Python 3.7. A model for NeatMS was generated from peaks generated from DS1 via XCMS3. In total 164 'High Quality', 57 'Low Quality', and 1497 'Noise' peaks have been labelled for training. We then trained the network viaTransfer Learning, using default parameters as provided in the NeatMS tutorial for all parameters. Afterwards, NeatMS was applied to label the same 9 aligned XCM3 outputs generated for the unsupervised clustering investigation, described above. Using R, different filters were applied to the NeatMS output so each NPP feature was required to contain a minimum of 0, 1, 3, 5, 8, or 10 'High Quality' peaks, while all other features were removed. For the extraction of NPP performance metrics via mzRAPP, BM1 was filtered so it contained only features which contained a peak for each of the 10 samples.

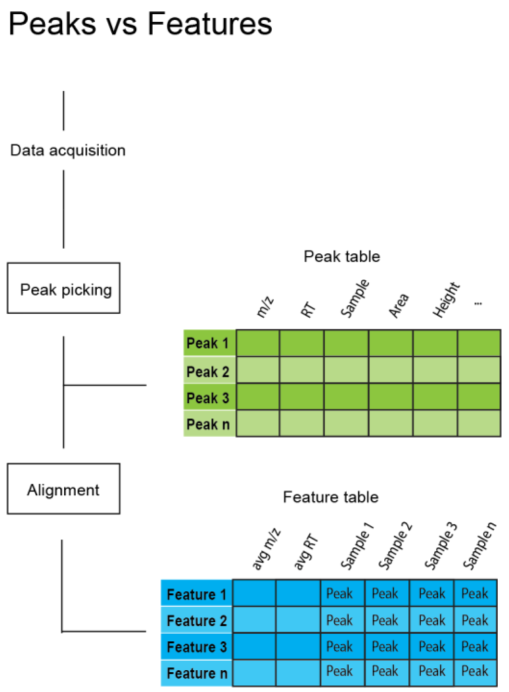


**Figure S1.** Major steps of NPP and glossary: Chromatographic peaks with an associated retention time and mz value are referred to as peaks. Multiple peaks aligned across several samples with an associated average mz and retention time value are referred to as features.

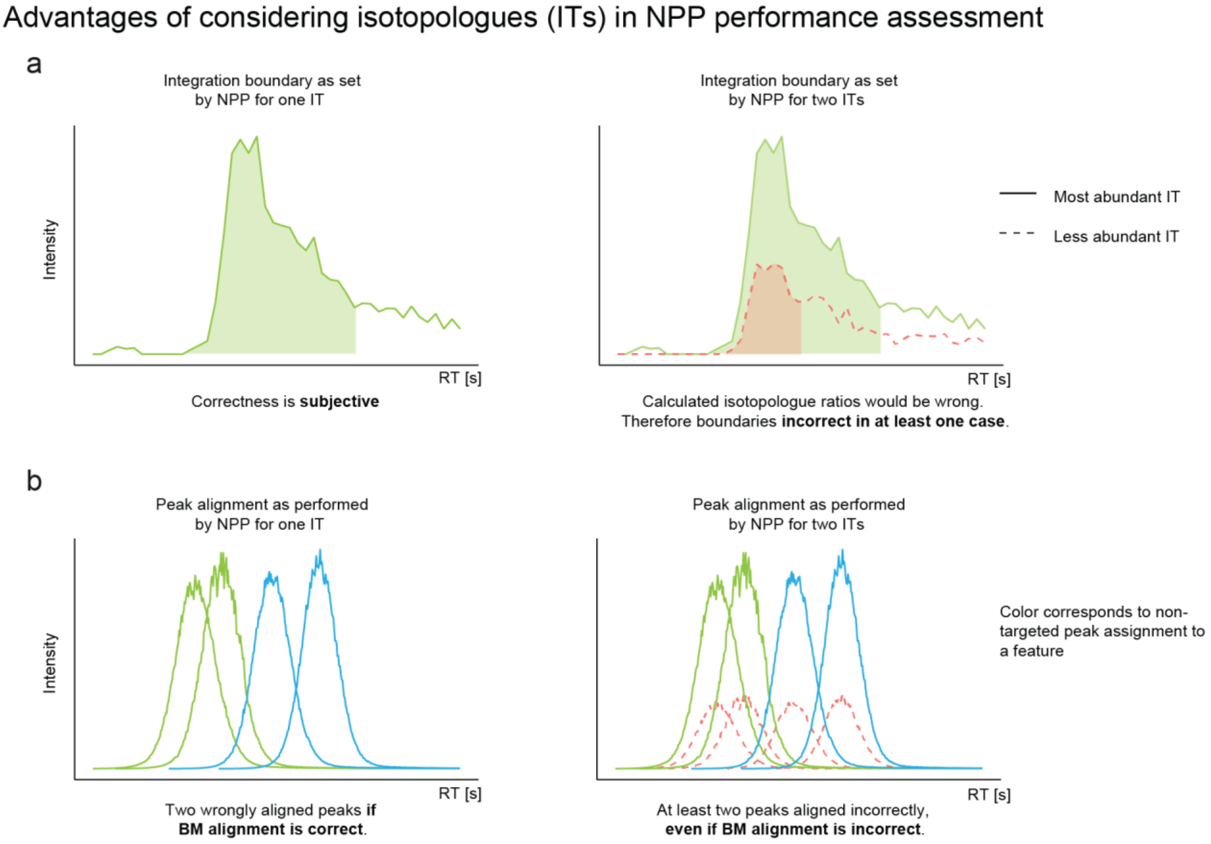


**Figure S2.** Considering isotopologues (ITs) for non-targeted data pre-processing (NPP) reliability assessment brings several key advantages. (a) By relying on IT ratios rather than exact benchmark (BM) peak areas, the evaluation of set integration boundaries becomes less subjective. (b) Considering the alignment of an additional IT feature shows that the same NPP algorithm assigned all of those into the same feature. Therefore at least two peaks must have been assigned incorrectly by the non-targeted algorithm, even when the BM alignment is incorrect.

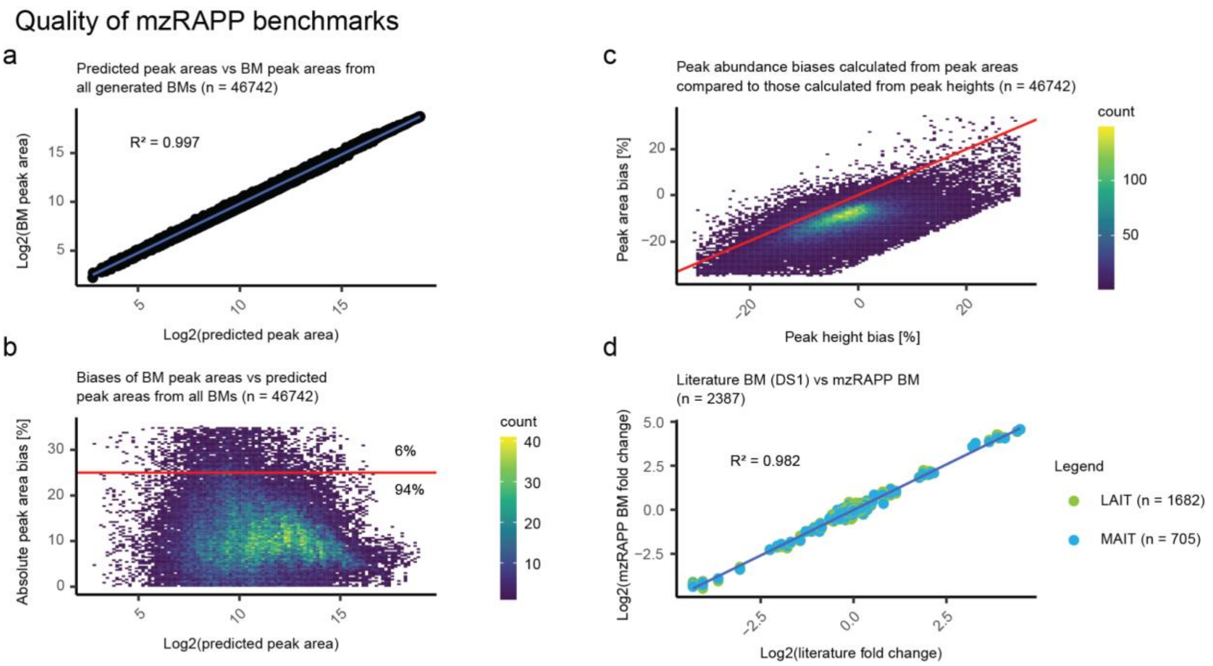


**Figure S3.** Plots characterizing benchmarks (BM) generated via mzRAPP. The peak area/height of low abundant isotopologues (LAITs) can be predicted from the peak area/height of the respective most abundant isotopologue (MAIT) when the molecular composition is known. (a) The plot demonstrates that this linear relationship is given for all peaks in mzRAPP generated BMs. (b) The plot shows how the relative bias of this prediction is below 25% for 94% of all BM peaks. (c) Small differences in biases calculated from peak heights to those from peak areas form an additional line of evidence for high-quality peak abundances. (d) Fold changes between two sample groups published for data set 1[9] were reproduced by mzRAPP not only for the reported MAIT, but also for LAITs underlining mzRAPPs ability to produce peaks with reliable abundances.

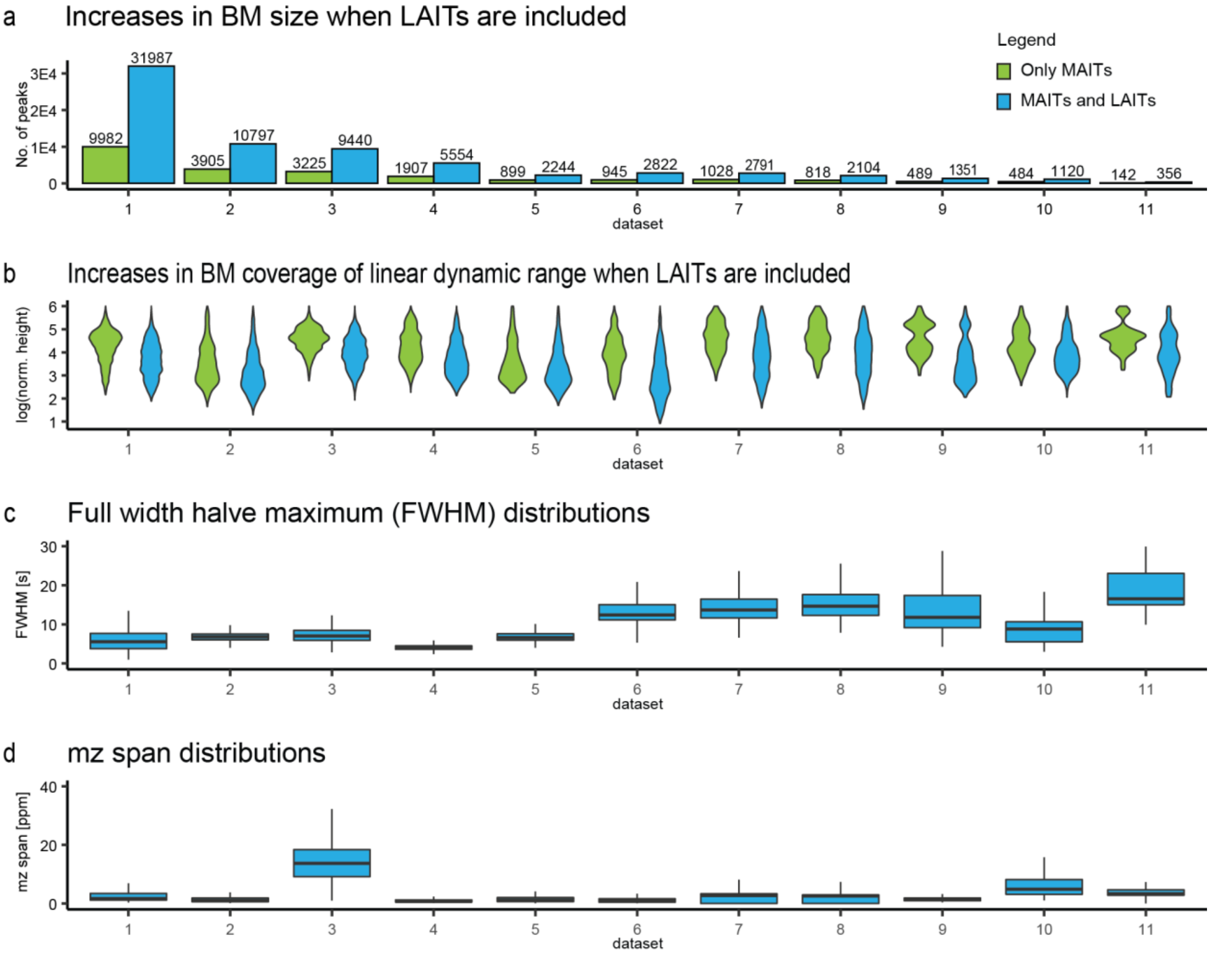


**Figure S4.** Eleven different benchmarks (BM) from different data sets (1-11, see supplement) have been produced using mzRAPP. (a) The bar plot shows the increased number of peaks achieved by adding peaks from all detectable isotopologues (blue) compared to only considering the most abundant isotopologue (MAIT). (b) The addition of lower abundant isotopologue (LAIT) peaks increased the benchmark coverage, as can be seen in the violin plots visualizing the linear dynamic range. In some cases, the covered linear dynamic range increased about two orders of magnitude. (c & d) Finally, the distribution of full width at half maximum (FWHM) and mz range is shown for all BM peaks.

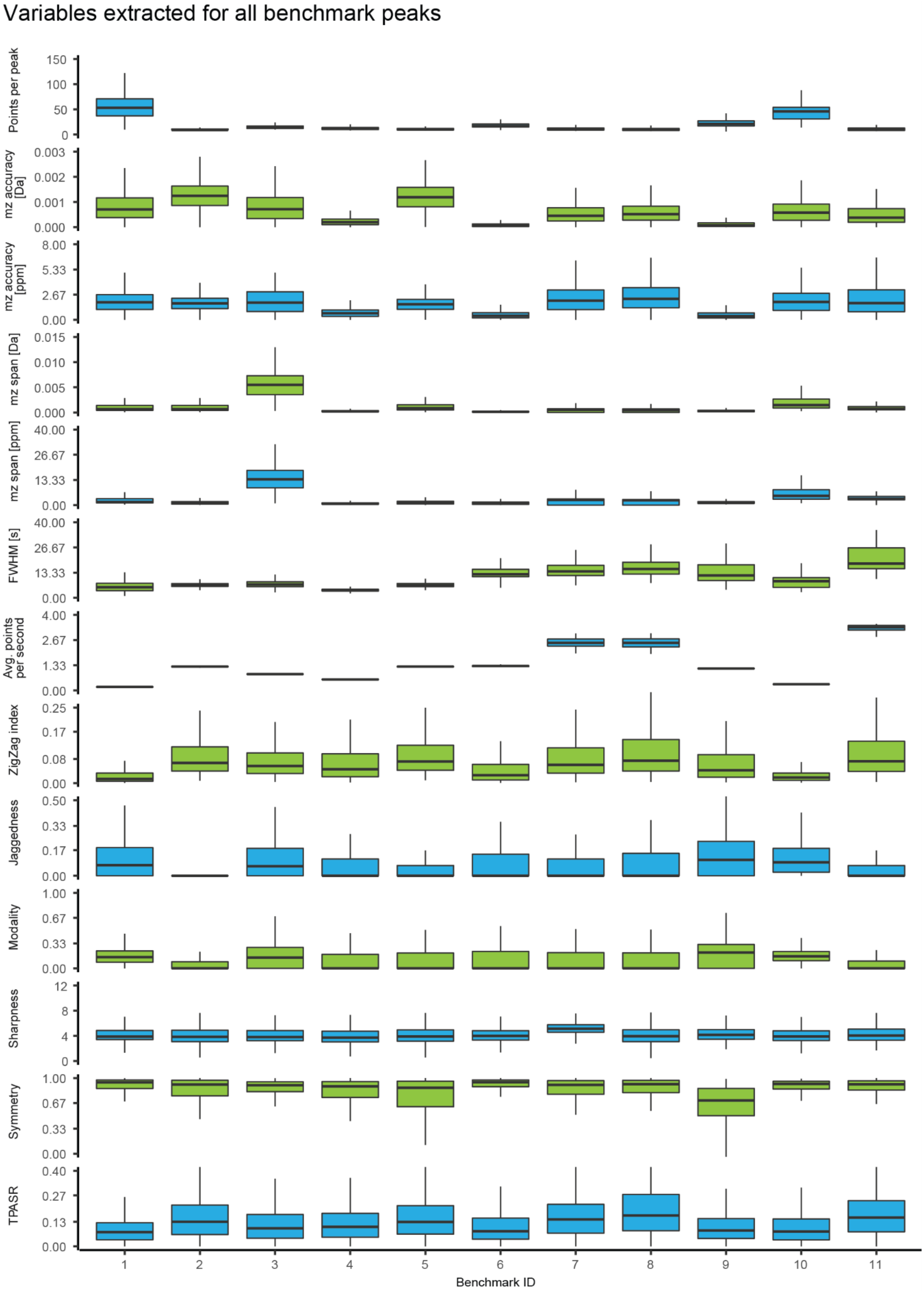


**Figure S5.** Distribution of peak variables for all generated benchmarks (outliers not shown).

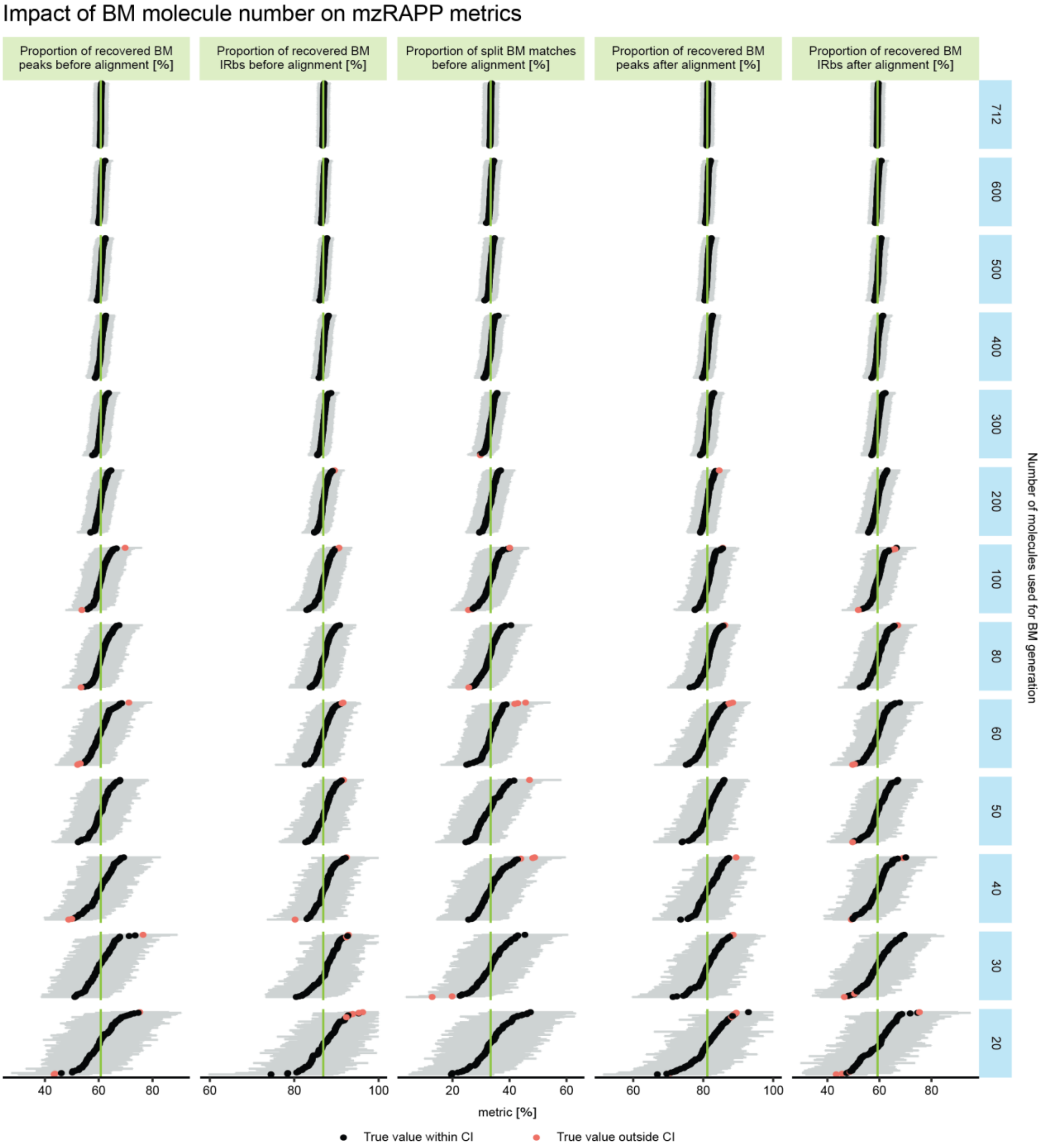


**Figure S6.** A benchmark was generated for dataset 1 considering 712 molecules und used for NPP assessmentStepwise reduction of molecules used for BM generation (molecules were sampled at random from all 712 molecules (n = 100) for molecule numbers ranging from 20 to 712) largely affected the size of confidence intervals (CI; as estimated by mzRAPP via bootstrapping with R = 1000 and confidence level = 0.99), while true metrics, as estimated from the original BM (green lines) stayed largely within within the reported CI of respective NPP assessments. (CI values above 100 were reduced to 100)

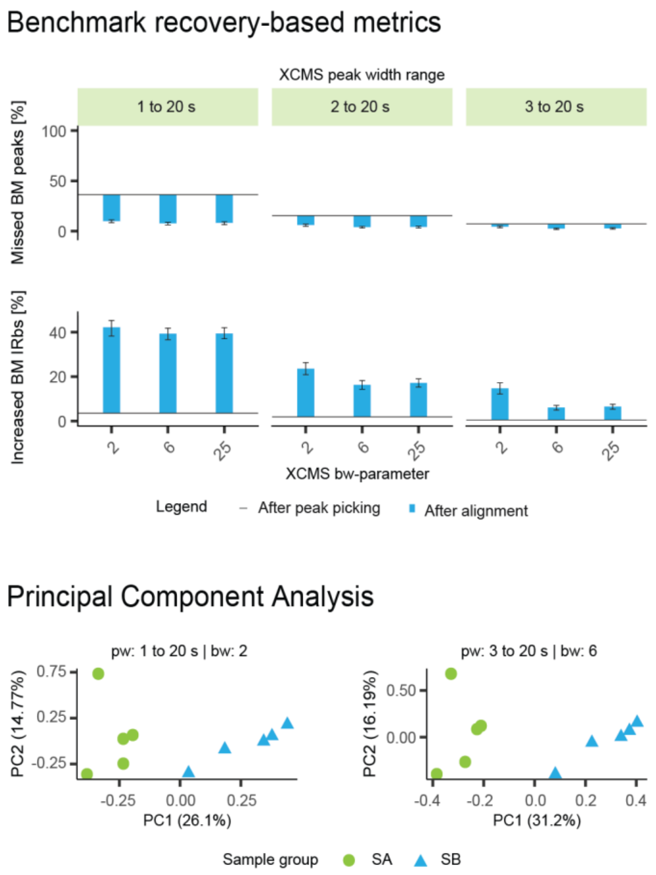


**Figure S7.** Benchmark (BM) recovery metrics are given for nine non-targeted data pre-processing (NPP) experiments done for dataset 1. Principal Component Analysis (PCAs) have been provided for two of them (see Figure PCAs_alternative (supplement) for the additional PCAs). As can be seen there is significant variability in the proportion of recovered isotopologue ratio biases (IRbs), while PCAs led to successful clustering of the introduced sample groups in all cases.

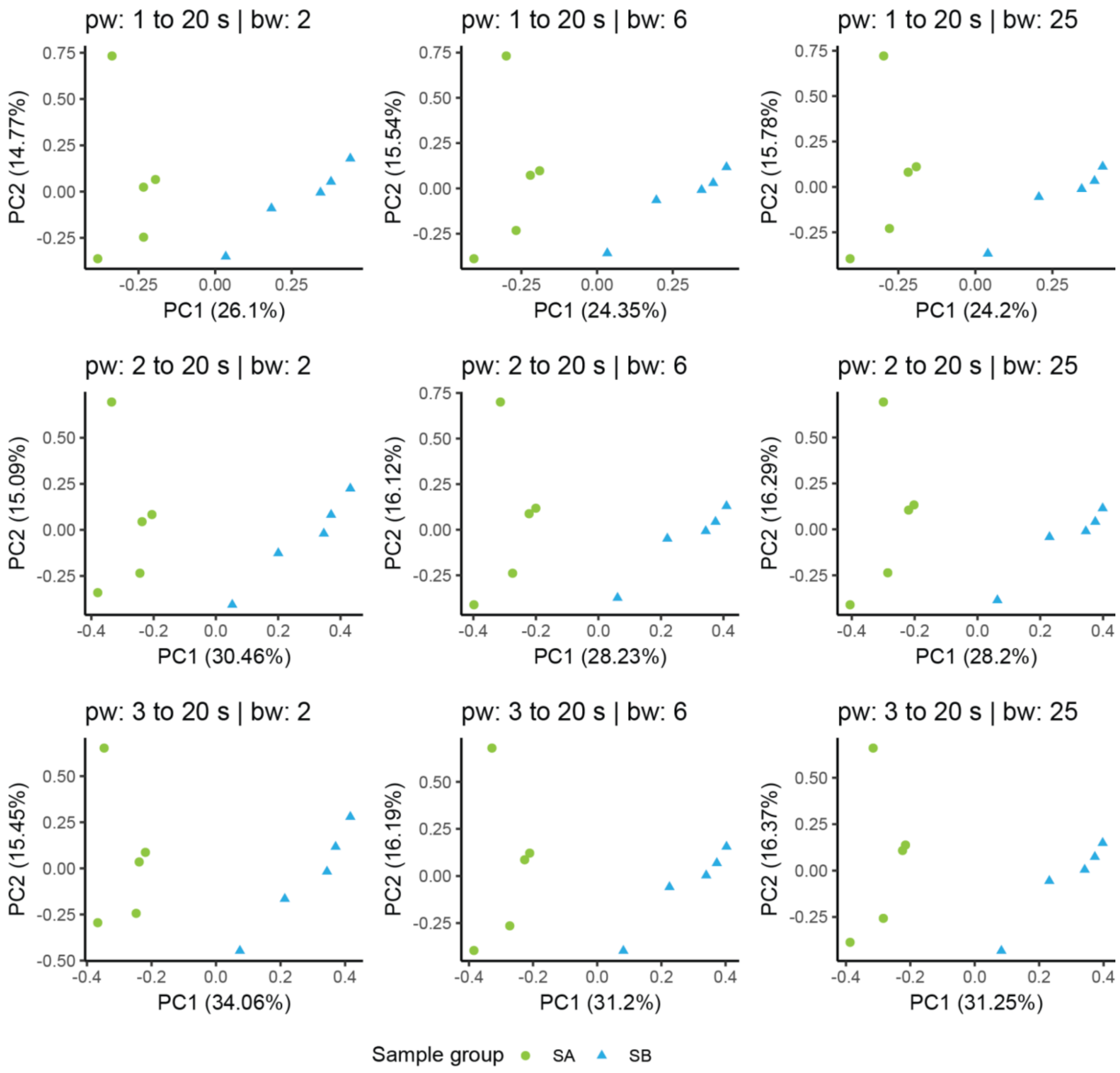


**Figure S8.** Principal Component Analysis were done for nine NPP experiments performed via XCMS3 on DS 1, with sample groups SA and SB. Non-targeted data pre-processing parameters were adapted in their minimum allowed peak width (pw) ranging from 1-20 s to 3-20 s and their bandwidth (bw), width value 2, 6, and 25.

**Table S1.** Overview of used datasets and generated benchmarks

| Dataset/Benchmark ID | Chromatography | Mass spectrometer | Polarity | Samples | Molecules | Peaks | Features | Reference |
|---|---|---|---|---|---|---|---|---|
| 1 | ZORBAX Eclipse C18 (RP) | Q Exactive HF | pos | 10 | 712 | 31987 | 3513 | [9] |
| 2 | ACQUITY HSS T3 (RP) | Q Exactive HF | pos | 59 | 104 | 10797 | 388 | MTBLS 1876 [28] |
| 3 | ZORBAX Eclipse C18 (RP) | Sciex TripleTOF6600 | pos | 8 | 451 | 9440 | 1725 | [9] |
| 4 | Kinetex EVO C18 (RP) | Q Exactive Plus | pos | 49 | 195 | 5554 | 919 | MTBLS 1455 [24] |
| 5 | ACQUITY HSS T3 (RP) | Q Exactive HF | pos | 9 | 119 | 2244 | 366 | MTBLS 1876 [28] |
| 6 | SeQuant ZIC pHILIC | Q Exactive Plus | pos | 26 | 77 | 2822 | 338 | MTBLS 706 [29] |
| 7 | SeQuant ZIC pHILIC | LTQ Orbitrap | pos | 30 | 42 | 2791 | 154 | MTBLS 265 [30] |
| 8 | SeQuant ZIC pHILIC | LTQ Orbitrap | pos | 30 | 36 | 2104 | 126 | MTBLS 267 [30] |
| 9 | SeQuant ZIC pHILIC | Q Exactive HF | neg | 24 | 26 | 1351 | 96 | MTBLS 2672 [31] |
| 10 | ACQUITY HSS T3 (RP) | Sciex X500R QTOF | pos | 15 | 44 | 1120 | 126 | MTBLS 1785 [22] |
| 11 | SeQuant ZIC pHILIC | LTQ Orbitrap | neg | 30 | 10 | 356 | 28 | MTBLS 265 [30] |

Reference IDs refer to the reference section in the main publication.
